# Root system architecture responses to high-temperature stress in synthetic-derived wheat lines reveal distinct adaptive patterns

**DOI:** 10.64898/2026.04.24.720666

**Authors:** Sultan Md Monwarul Islam, Izzat Sidahmed Ali Tahir, Kinya Akashi

## Abstract

High-temperature stress poses a major threat to wheat productivity, particularly during early developmental stages. Root system architecture (RSA) plays a key role in stress adaptation; however, its variation under high-temperature stress remains insufficiently characterized, especially in genetically diverse populations. In this study, we evaluated RSA responses of representative genotypes from a Multiple Synthetic Derivatives (MSD) wheat population under control and high-temperature conditions using a time-resolved two-dimensional phenotyping platform. High-temperature stress significantly affected most root traits, with lateral root–related parameters, including second pair seminal root length (SPSRL), root system width (RSW), and convex hull area (CHA), showing relatively greater responsiveness than vertical traits. Integrative analyses combining stress indices and multivariate approaches revealed distinct genotypic response patterns. MSD417 and MSD034 maintained higher root performance under stress, indicating greater tolerance, whereas MSD392 exhibited pronounced sensitivity, and MSD054 showed limited responsiveness. These findings suggest the importance of distinguishing between active stress tolerance and apparent stability and indicate that lateral root–related traits may represent useful targets for selection. Overall, the findings of this study validate the practical usefulness of the RSA screening approach and identify MSD genetic resources harboring RSA traits relevant to breeding heat-resilient wheat.

## 1. Introduction

Wheat (*Triticum aestivum* L.) is the most widely cultivated cereal crop worldwide and provides a major source of calories for more than 35% of the global population **[1]**. With the global population projected to reach 9.6 billion by 2050, a substantial increase in wheat production will be required to meet future food demand **[2]**. However, climate change, particularly the rise in global temperatures, poses a significant threat to wheat productivity **[3]**. It has been estimated that wheat yield decline by 4.1–6.4% for every 1 °C increase in temperature **[4,5]**. Wheat growth is generally favored within a moderate thermal range (approximately 22/14 °C day/night), although the crop can perform well across a wider range of conditions. In addition, optimal seedling vigor has been reported around 25 °C under controlled conditions **[6]**. Nevertheless, exposure to temperatures above these favorable ranges, particularly during sensitive growth stages, can adversely affect physiological processes and productivity **[7,6,8,9]**. Elevated root-zone temperature, either alone or in combination with elevated shoot temperature, significantly reduced wheat growth, photosynthetic performance, and grain yield at various developmental stages **[10,11]**.

Adaptation to heat stress involves multiple physiological and morphological traits. While breeding efforts have focused on above-ground characteristics, increasing evidence suggests that below-ground traits also contribute substantially to heat resilience. Root system architecture (RSA), defined by traits such as root length, number, angle, and spatial distribution, plays a critical role in plant growth by regulating water and nutrient acquisition and mediating adaptation to environmental conditions **[12,13]**. RSA has been proposed as a key target for breeding climate-resilient crops, particularly due to its plasticity in response to environmental stresses **[14–16]**. Early-stage root development is especially important, as it can influence subsequent root system formation and overall plant performance **[17]**. Despite its importance, RSA remains underutilized in wheat breeding programs, largely due to the difficulty of root phenotyping. Evaluating RSA under field conditions remains challenging due to limited accessibility, destructive sampling requirements, and low temporal resolution, making it difficult to capture dynamic root responses **[12,18–21]**. Although various root phenotyping platforms have been developed **[22–31]**, continuous, high-resolution monitoring of RSA dynamics remains a major challenge. To address this bottleneck, a practical two-dimensional RSA phenotyping platform has recently been developed. However, its validation has so far been limited to a single multiple synthetic derivative (MSD) line, which restricts confidence in its broader applicability **[32]**.

In parallel, progress in improving RSA is limited by the narrow genetic diversity of modern bread wheat, particularly in the D genome. *Aegilops tauschii*, the D-genome donor of bread wheat, is a valuable source of novel alleles for stress adaptation. The Multiple Synthetic Derivative (MSD) wheat population represents a powerful platform for exploring genetic variation derived from *Ae. tauschii*. This population was developed through repeated crossing and backcrossing of 43 synthetic wheat lines with the common wheat cultivar Norin 61 **[33,34]**. By incorporating genomic contributions from diverse *Ae. tauschii* accessions, the MSD population captures a broad spectrum of genetic diversity. The D genome has been widely recognized as an important source of allelic variation associated with abiotic stress adaptation, including tolerance to heat and drought, and contributes to traits related to resource acquisition **[33–36]**. In addition to general stress responses, variation derived from *Ae. tauschii* has been associated with root system traits and nutrient uptake, suggesting its potential role in shaping RSA under stress conditions **[35,37–39]**. Therefore, the genetic contribution of the D genome in the MSD population provides an opportunity to investigate diversity in root system architecture.

Based on field performance in Sudan, one of the hottest wheat-growing regions in the world, several MSD genotypes (MSD034, MSD054, MSD296, MSD392, and MSD417) were identified as heat-tolerant candidates, with accompanying physiological and molecular analyses revealing distinct stress-response patterns among these genotypes **[40,41]**. However, apart from MSD417, the RSA responses of the selected MSD lines have not been systematically characterized. Therefore, the objectives of this study were to (i) further validate a recently developed RSA phenotyping method by expanding its evaluation across multiple genotypes under control and high-temperature conditions, and (ii) characterize RSA variation and identify promising root traits potentially associated with heat stress adaptation. The findings of this study are expected to strengthen the methodological basis for RSA phenotyping and support the utilization of MSD germplasm in breeding programs targeting wheat heat resilience.

## 2. Results

### 2.1. Comparison of the trait parameters

Root and shoot traits of various MSD genotypes and their recurrent parent N61 were compared at control temperatures (22 °C/18 °C day/night) and at high temperatures (42 °C/18 °C) during the second half of the eight-day growth period. To quantitatively evaluate root system characteristics, the trait parameters were measured as defined in Table 1. Analysis of variance (ANOVA) revealed that genotypes (G), contrasting temperature conditions (Environment, E), and their interaction (G × E) significantly influenced most root and shoot traits (Table 1). Highly significant genotypic differences (*p* ≤ 0.001) were observed for nearly all traits, including total root length (TRL), root system width (RSW), and convex hull area (CHA), indicating substantial genetic variability within the selected lines. Notably, mean square values for E were markedly higher than those for G across most traits, suggesting that high temperature was the primary driver of phenotypic variation. While E significantly impacted nearly all traits (*p* ≤ 0.001), shoot dry weight (SDW) remained notably stable across treatments. Significant G × E interactions for traits such as root dry weight (RDW) and CHA (*p* ≤ 0.01) indicate differential genotypic plasticity in root biomass allocation under high temperature stress. In contrast, the lack of significant G × E interaction for rooting depth (RD) and specific root length (SRL) suggests these traits were more genetically stable across environments.

**Table 1.** Analysis of variance for different root and shoot traits under high temperature conditions compared to control conditions in wheat genotype.

| Traits | Abbreviation | Unit | Mean square (MS) |  |  |  |
| --- | --- | --- | --- | --- | --- | --- |
|  |  |  | Genotype (G) | Environment (E) | G × E interaction | Residual |
| Rooting depth | RD | cm | 9.9 ns | 615.32 *** | 0.97 ns | 7.18 |
| Shoot height | SH | cm | 21.11 *** | 38.38 *** | 1.99 ns | 1.31 |
| Root dry weight | RDW | mg | 11.1 *** | 94.81 *** | 5.38 ** | 1.16 |
| Shoot dry weight | SDW | mg | 43.34 *** | 3.89 ns | 5.86 * | 2.35 |
| Root to shoot ratio | RSR |  | 0.13 ** | 0.14 ** | 0.01 * | <0.01 |
| Total root length | TRL | cm | 1014.68 *** | 16384.7 *** | 28.73 ns | 66.82 |
| Specific root length | SRL | cm mg<sup>-1</sup> | 2.64 *** | 26.18 *** | 0.69 ns | 0.40 |
| Primary root length | PRL | cm | 65.79 *** | 796.5 *** | 5.99 ns | 13.56 |
| 1<sup>st</sup> pair seminal root length | FPSRL | cm | 26.25 *** | 672.15 *** | 10.81 * | 3.99 |
| 2<sup>nd</sup> pair seminal root length | SPSRL | cm | 102.02 *** | 161.15 *** | 7.32 ns | 6.72 |
| Root system width | RSW | cm | 177.22 *** | 529.24 *** | 47.2 * | 18.09 |
| Convex hull area | CHA | cm<sup>2</sup> | 56761.94 *** | 403428.16 *** | 13979.88 ** | 3138.52 |
| 1<sup>st</sup> pair seminal root angle | FPSRA | degree | 2197.34 *** | n.a. | n.a. | 217.51 |
| 2<sup>nd</sup> pair seminal root angle | SPSRA | degree | 3326.07 *** | n.a. | n.a. | 285.51 |
| df |  |  | 5 | 1 | 5 | 60 |

\* $p < 0.05$ , \*\* $p < 0.01$ , \*\*\* $p < 0.001$ , ns = not significant, n.a. = not applicable as both pair of seminal roots developed before imposing high temperature stress and had no treatment effect on seminal root angle development

Figure 1 presents genotype-wise comparisons of root architectural traits under control and high-temperature conditions. Among these traits, RD, a key proxy for water and nutrient acquisition from deep soil layers, was significantly reduced across all tested genotypes, with reductions ranging from 20.76% to 27.35% (Figure 1A and Supplementary Table S1). RD showed no significant genotypic differences in both control and high-temperature conditions. In contrast, the response of shoot height (SH) to high-temperature stress varied among genotypes (Figure 1B). High-temperature stress led to significant reductions in height in N61 and MSD392, with MSD392 exhibiting the greatest decrease (16.68%). Contrastingly, SH remained relatively stable in MSD034, MSD054, MSD296, and MSD417. Under control conditions, MSD417 and MSD296 exhibited the highest root dry weight (RDW) and shoot dry weight (SDW), reflecting high growth potential (Figure 1C, D). High-temperature stress significantly reduced RDW in MSD034, MSD296, MSD392, and MSD417. In contrast, RDW remained stable in N61 and MSD054, indicating that root biomass allocation was maintained under stress (Figure 1C and Supplementary Table S1). Although mean SDW values tended to be lower under high-temperature stress across several genotypes, these differences were not statistically significant, except for MSD417 (Figure 1D). Biomass partitioning, expressed as the root-to-shoot ratio (RSR), varied significantly among genotypes. Under control conditions, MSD034 exhibited the highest RSR (1.02 ± 0.10), while other genotypes ranged from 0.68 ± 0.03 to 0.87 ± 0.10 (Figure 1E and Supplementary Table S1). High-temperature stress significantly reduced RSR in MSD034, MSD054, and MSD392, whereas RSR changes were not significant in the other genotypes. These observations suggest that root development was more sensitive to high-temperature stress than shoot growth in the affected genotypes.

**Figure 1.**
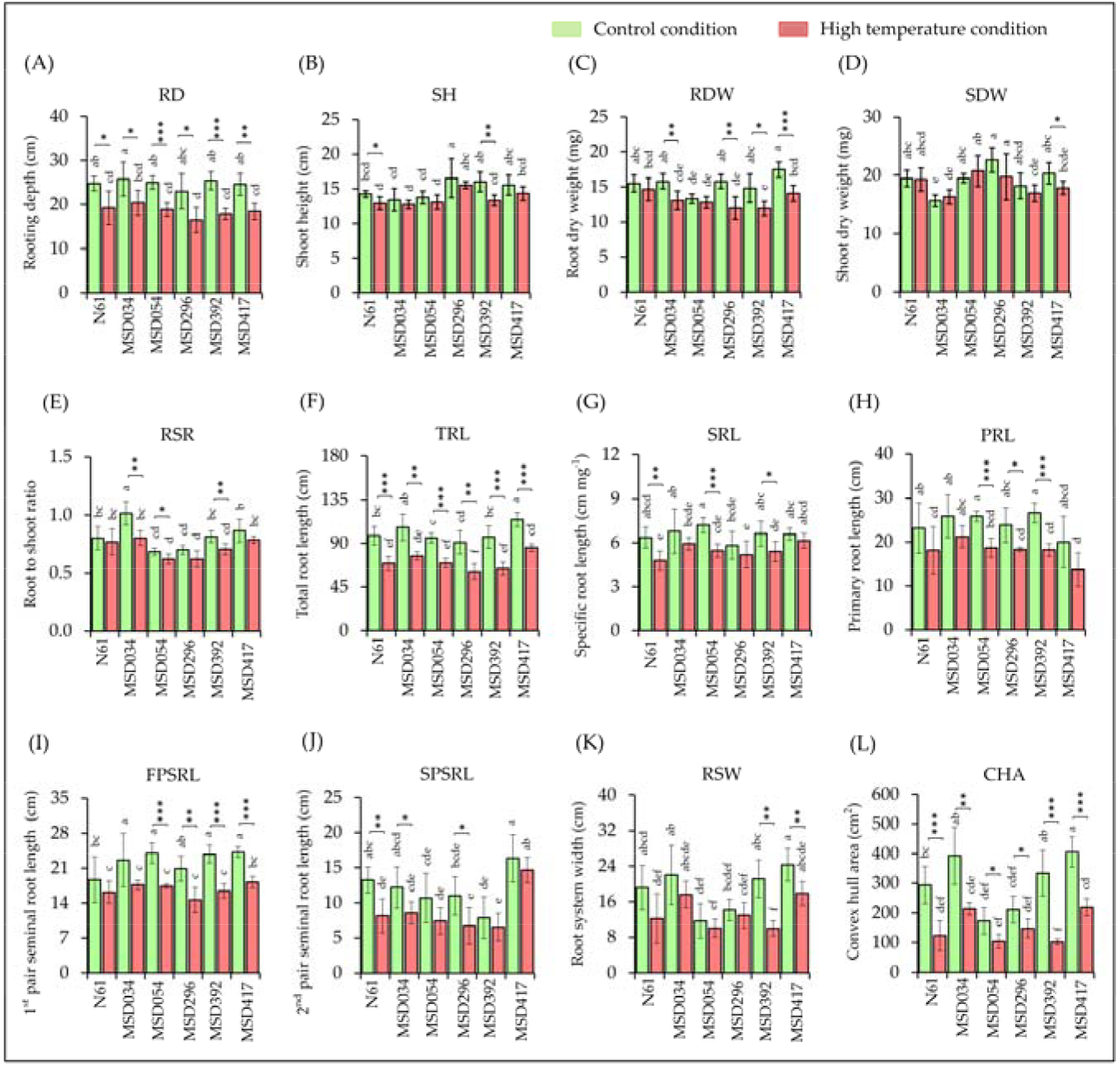
Variation in root and shoot morphological traits in contrasting temperatures of different wheat genotypes at day eight after germination. Genotype data were compared using Tukey’s HSD post hoc test, and different letters indicate significant differences among genotypes under contrasting temperature conditions. Data for each genotype at contrasting temperatures were compared using Student’s t-test (*p < 0.05, **p < 0.01, ***p < 0.001), n = 6. Please see Table 1 for trait abbreviations.

To evaluate the geometric configuration and soil coverage potential of the root system, total root length (TRL), specific root length (SRL), and the lengths of major roots were analyzed (Figure 1F–J, Supplementary Table S1). Under control conditions, TRL of the tested genotypes ranged from 91.14 ± 12.01 to 114.56 ± 7.20 cm, where MSD417 had the highest mean value (Figure 1F and Supplementary Table S1). High TRL value of MSD417 has been reported previously [32]. High-temperature stress significantly reduced TRL across all genotypes, and MSD417 maintained the highest mean values with the smallest reduction (25.18%), whereas MSD392 exhibited the greatest sensitivity, with a 33.76% reduction. Under control conditions, mean SRL values, an indicator of root fineness and resource acquisition efficiency, ranged from 5.81 ± 0.94 to 7.20 ± 0.53 cm mg^-1^. MSD054 exhibited the highest value, indicating a strategy of producing greater root length per unit root biomass (Figure 1G, Supplementary Table S1). Under high-temperature stress, N61, MSD054, and MSD392 showed significant reductions in SRL, whereas MSD034, MSD296, and MSD417 showed no statistically significant changes. MSD417 exhibited the smallest mean reduction (6.78%), whereas the remaining genotypes showed mean reductions ranging from 14.70% to 24.83% (Supplementary Table S1). Although primary root length (PRL) showed little significant genotypic differences in the control conditions, considerable genotypic variations were observed in the 1^st^ pair seminal root length (FPSRL) and 2^nd^ pair seminal root length (SPSRL), particularly in MSD417 for high SPSRL value (Figure 1H–J, Supplementary Table S1). FPSRL was highly sensitive to high temperatures in MSD054, MSD296, MSD392, and MSD417, whereas no statistically significant changes were detected in N61 and MSD034, despite lower mean values under stress. Similarly, SPSRL was significantly reduced in N61, MSD034, and MSD296, whereas changes in the remaining three genotypes were not statistically significant. MSD417 maintained the highest absolute SPSRL even under stress, with the smallest percent reduction (10.38%), despite significant reductions in TRL and FPSRL, suggesting prioritized maintenance of certain root types in this genotype.

Seminal root angles were formed during the early stages of seedling development, prior to high-temperature treatment, and differed markedly among genotypes (Figure 2). MSD417 exhibited the widest angles for both the 1^st^ and 2^nd^ pairs of seminal roots (FPSRA and SPSRA, respectively). MSD034 and MSD296 also had a wide SPSRA, comparable to MSD417, suggesting that these genotypes possess a naturally broader basal footprint than the other genotypes, which had more vertically oriented root systems. Under control conditions, the mean root system width (RSW), which is influenced by the angle and length of the second pair of seminal roots (SPSRA and SPSRL, respectively), was greatest in MSD417, followed by MSD034 and MSD392 (Figure 1K, Supplementary Table S1). High-temperature stress significantly reduced RSW in MSD392 and MSD417; however, MSD417 and MSD034 retained higher RSW values than the other genotypes. Convex hull area (CHA), which reflects total soil area explored by roots, varied significantly among genotypes. Under control conditions, MSD417 and MSD034 exhibited higher mean CHA values than the other genotypes (Figure 1L and Supplementary Table S1). High-temperature stress significantly reduced CHA in all genotypes, with percentage reductions ranging from 30.08% to 58.04%. Once again, MSD417 and MSD034 retained higher mean CHA values than the other genotypes.

**Figure 2.**
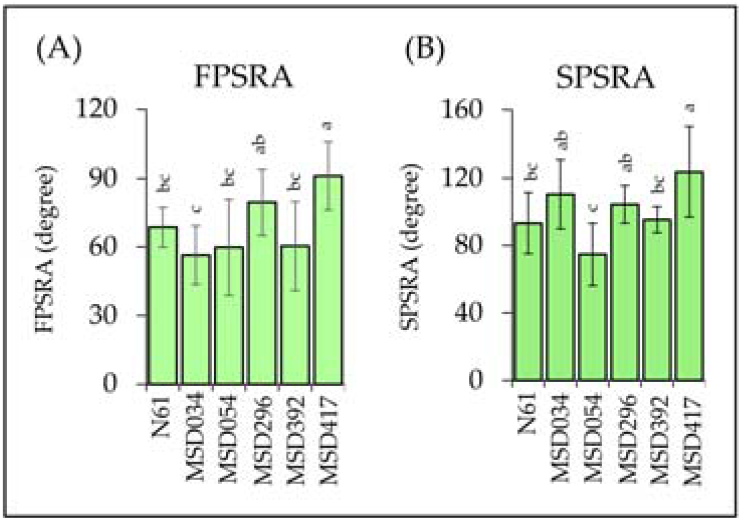
Variation in seminal roots angle development of different wheat genotypes at day eight after germination. Data for the genotypes were compared with Tukey’s HSD post hoc test, and different letters denote significant differences among the genotypes. n = 12. Please see Table 1 for trait abbreviations.

### 2.2. Hierarchical cluster analysis and distance matrix of the traits

To evaluate the relationships between genotypes and their morphological responses to high-temperature stress, hierarchical cluster analysis (HCA) and a pairwise distance matrix were performed (Figure 3). The HCA dendrogram and heatmap partitioned the genotypes into distinct clusters based on their phenotypic profiles under control and high-temperature conditions (Figure 3A). The most significant result was the clear separation between control and high-temperature-stressed plants. The control group exhibited higher values for biomass and architectural traits, including RDW, TRL, RSW, and CHA. In contrast, the high-temperature clade was characterized by widespread reductions in root traits, with N61, MSD054, MSD296, and MSD392 showing the most pronounced declines in RD, TRL, SPSRL, and CHA. The horizontal dendrogram at the top revealed characteristic trait correlations, including a close association between RSW and CHA, indicating that these architectural parameters respond similarly to high-temperature stress.

**Figure 3.**
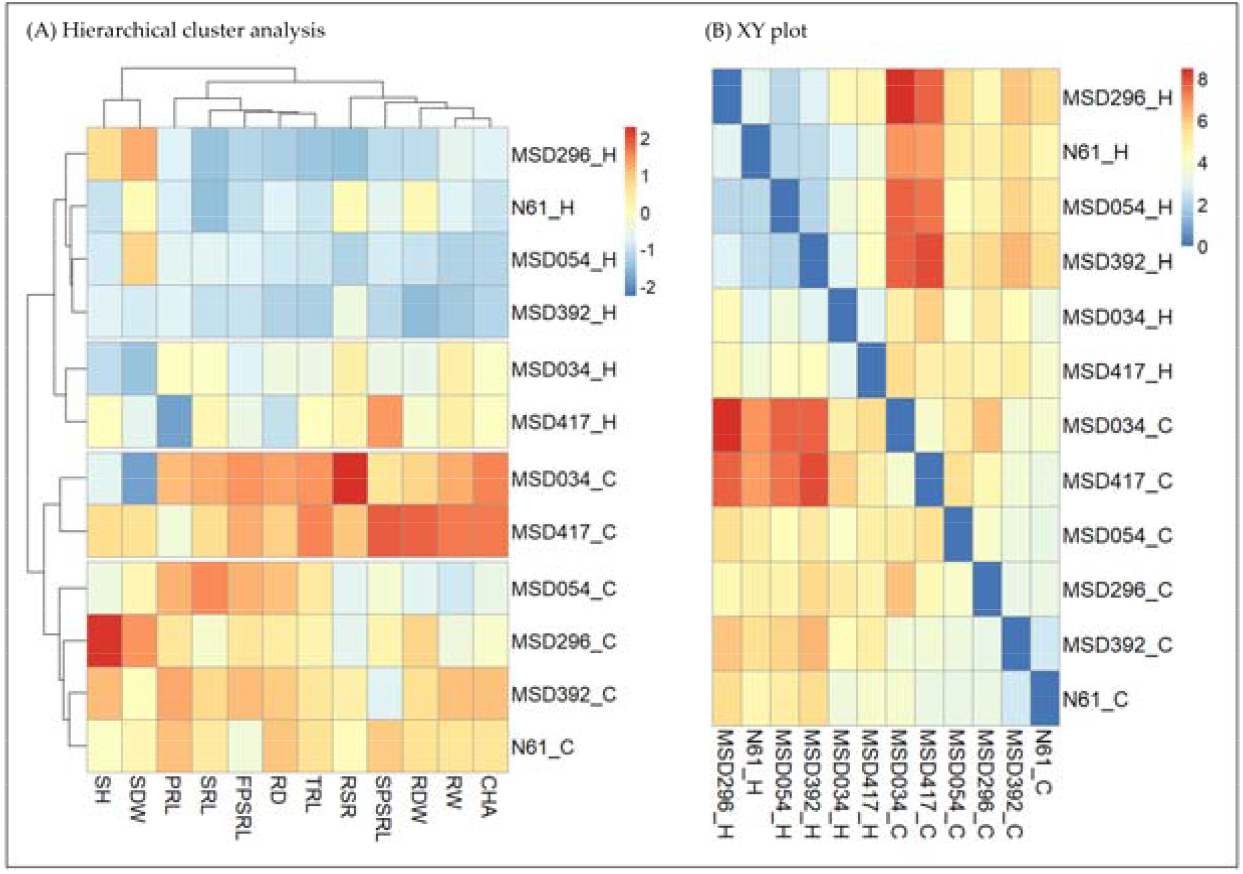
Hierarchical cluster analysis and phenotypic distance of wheat genotypes under control and heat stress conditions. (A) Double dendrogram and heatmap visualizing the clustering of wheat genotypes based on measured root and shoot traits. Rows represent genotypes in contrasting temperatures, and columns represent the measured traits. The color scale indicates standardized values (Z-scores), with red representing values above the mean and blue representing values below the mean. The clustering was performed using Ward’s method based on Euclidean distances. (B) Distance matrix illustrating the phenotypic dissimilarity between samples. The color gradient ranges from blue (low distance, high similarity) to red (high distance, low similarity). Genotype name with _C extension explains plants in the control conditions, and genotype name with _H extension explains plants in the high-temperature conditions, n= 6. Please see Table 1 for trait abbreviations.

The heatmap of pairwise phenotypic distances revealed large distances between control samples of MSD034 and MSD417 and the high-temperature-stressed samples of MSD296, N61, MSD054, and MSD392, indicating pronounced phenotypic divergence between the high-performing genotypes under control conditions and the other genotypes under high-temperature conditions (Figure 3B). Under control conditions, N61 and MSD392 showed lower distance values, suggesting similar baseline growth strategies. Interestingly, several genotypes (MSD296, N61, MSD054, and MSD392) showed reduced pairwise distances under high-temperature stress compared with control conditions, forming a region of higher similarity in the distance matrix. This suggests that high-temperature stress acts as a physiological bottleneck, forcing different genotypes toward a similarly stunted phenotype.

### 2.3. Comparative evaluation of stress indices and genotypic stability

To enable an integrated comparison of genotypic performance under high-temperature stress, several stress indices were calculated from trait performance under control and stress conditions. Percent reduction (PR), stress susceptibility percentage index (SSPI), mean relative performance (MRP), and stress tolerance index (STI) were computed for all traits (Supplementary Tables S2–S5), and the resulting profiles were visualized using radar plots (Figure 4).

**Figure 4.**
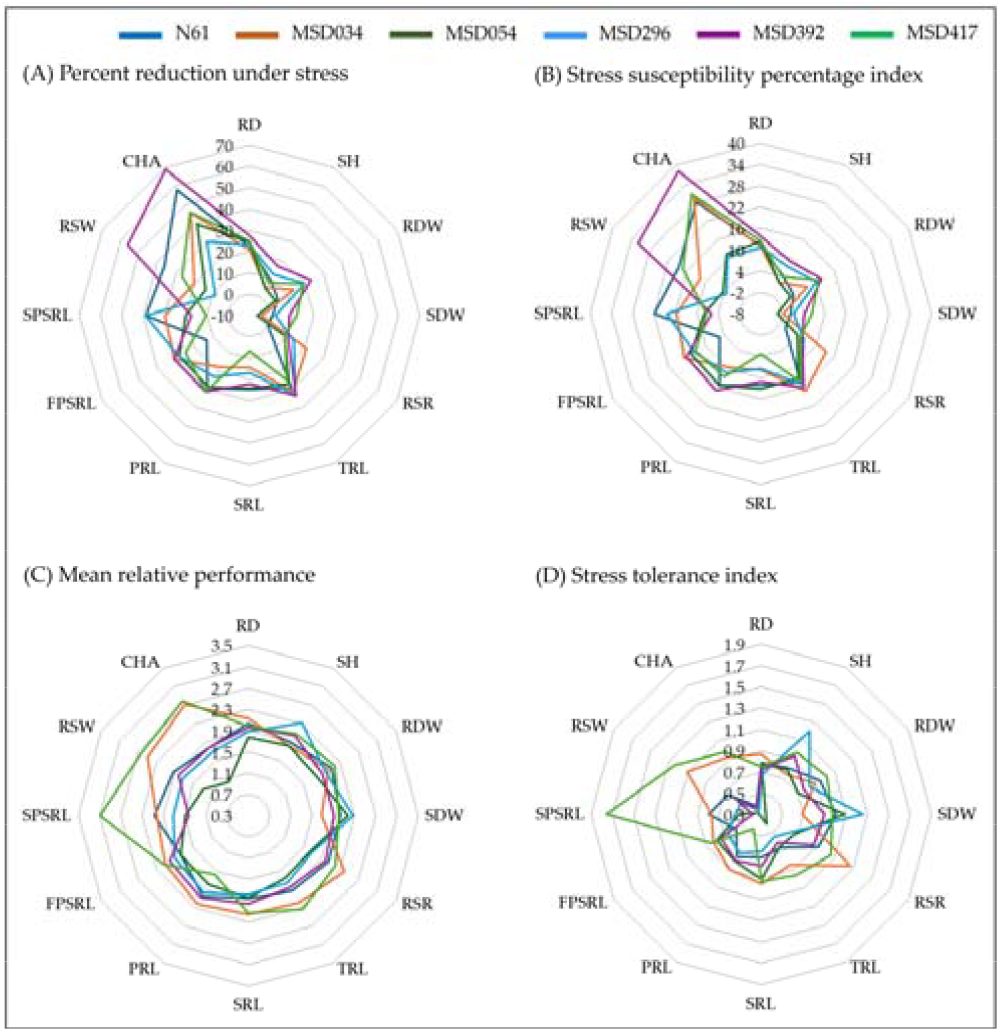
The radar plots showing the stress tolerance indices of different root and shoot traits of different genotypes due to high temperature stress. (A) Percent reduction of different traits, (B) Stress susceptibility percentage index, (C) Mean relative performance, and (D) Stress tolerance index. Please see Table 1 for trait abbreviations.

PR analysis identified convex hull area (CHA), root system width (RSW), and 2^nd^ pair seminal root length (SPSRL) as the traits most severely affected by high-temperature (Figure 4A, Supplementary Table S2). The largest reductions were observed in MSD392 (69.22% for CHA and 56.29% for RSW) and in N61 for SPSRL (38.56%). Across genotypes, MSD392 showed consistently high reduction in root traits, including RD (27.35%) and TRL (33.76%). In contrast, SDW was the most stable trait, with minimal reduction in N61 (0.93%) and slight increases in MSD054 and MSD034, as indicated by negative PR values (-6.51% and -4.35%, respectively). Overall, N61, MSD034, and MSD417 exhibited relatively greater stability, maintaining lower reductions across multiple root traits.

SSPI was used to quantify relative declines in performance, with lower values indicating greater stability (Figure 4B, Supplementary Table S3). Overall, most genotypes exhibited moderate susceptibility across root length traits (RD, TRL, and PRL). Notably, MSD392 showed pronounced sensitivity in root expansion traits, particularly CHA (38.31) and RSW (31.75). In contrast, MSD054 exhibited the lowest overall SSPI (mean = 8.06), indicating greater stability. N61, MSD034, and MSD417 showed comparable performance, with mean SSPI values ranging from 11.18 to 11.44.

Mean relative performance (MRP) values indicated that certain genotypes performed better under high-temperature stress (Figure 4C, Supplementary Table S4). MSD417 showed the highest overall performance (mean MRP = 2.26), followed by MSD034 (2.17). The high performance of MSD417 was associated with certain root traits, with the highest values observed for SPSRL (3.09), CHA (2.79), and RSW (2.64). MSD034 exhibited consistently high values across traits, including RSR (2.39) and RSW (2.50), indicating stable performance. In contrast, MSD296 showed relatively high values in shoot traits, particularly SH (2.31) and SDW (2.27). MSD054 exhibited the lowest values across most traits.

The stress tolerance index (STI) further revealed patterns of genotypic variation under high-temperature stress (Figure 4D; Supplementary Table S5). MSD417 exhibited the highest overall tolerance (mean STI = 0.99), with elevated values in SPSRL (1.76) and RSW (1.24). MSD034 followed (mean STI = 0.90), showing high values in RSR (1.27) and RSW (1.11). In contrast, MSD054 exhibited the lowest STI values (mean = 0.68), reflecting reduced performance, particularly in CHA (0.20) and RSW (0.33).

### 2.4. Principal component analysis of genotype performance under high temperature stress

To comprehensively characterize genotypic responses to high-temperature stress, two complementary principal component analyses (PCAs) were performed: one based on raw phenotypic traits (Figure 5A) and the other based on stress indices (Figure 5B). In Figure 5A, the first two principal components explained 78% of the total phenotypic variation, with Dim1 and Dim2 accounting for 61% and 17%, respectively. Dim1 clearly separated genotypes by environmental conditions: control conditions (green circles) were distributed on the positive side, whereas high-temperature stress samples (red triangles) were located on the negative side. This separation was mainly associated with root architectural traits, including total root length (TRL), convex hull area (CHA), rooting depth (RD), and root system width (RSW), which were strong positive loadings on Dim1. In contrast, shoot-related traits such as shoot dry weight (SDW) and shoot height (SH) contributed more significantly to Dim2, suggesting a distinct axis of variation from root architecture. Genotypic differences were also evident within each condition. Under control conditions, MSD417 was closely associated with higher TRL and CHA, whereas MSD034 was characterized by higher root-to-shoot ratios (RSR). Under high-temperature stress, all genotypes shifted toward negative values along Dim1; however, MSD417 and MSD034 remained closer to the origin than the other genotypes, indicating a relatively greater maintenance of root architectur**a**l traits compared with genotypes such as MSD296 and MSD392.

**Figure 5.**
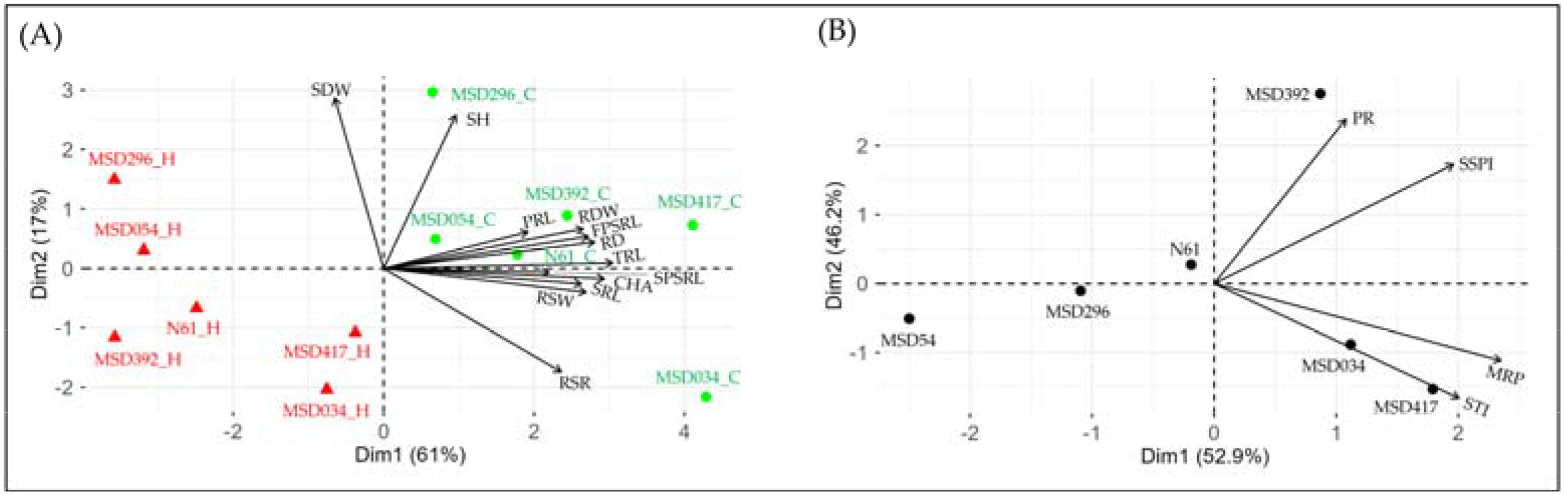
Principal Component Analysis (PCA) biplot of wheat genotypes under contrasting temperature conditions (A), and genotypic variation and clustering based on high temperature stress indices (B). Vectors indicate the direction and magnitude of trait associations. Genotypes oriented along specific vectors exhibit higher values for those traits. Genotype name with _C extension explains plants in the control conditions, and genotype name with _H extension explains plants in the high-temperature conditions. Please see Table 1 for trait abbreviations. PR = percent reduction, SSPI = stress susceptibility percentage index, RP = relative performance, and STI = stress tolerance index.

The stress index-based PCA (Figure 5B) provided another integrated representation of genotypic responses by combining multiple stress-related indices. The first two principal components explained a substantial proportion of the variation (Dim1 = 52.9%, Dim2 = 46.2%). In this biplot, performance- and tolerance-related indices, such as MRP and STI, were associated with the positive side of Dim1 and the negative side of Dim2, whereas stress sensitivity metrics, such as PR and SSPI, were located on the positive side of both Dim1 and Dim2. Genotypes were clearly differentiated along these axes. MSD417 and MSD034 were closely associated with STI and MRP, indicating higher overall performance and tolerance under high-temperature stress. In contrast, MSD054 was positioned on the negative side of Dim1, reflecting lower overall performance. MSD392 was separated, along with a higher Dim2 value, corresponding to elevated PR and SSPI values, indicating greater stress susceptibility. N61 and MSD296 were situated at intermediate positions, suggesting moderate responses across indices.

## 3. Discussion

Root system architecture (RSA) is a critical target for crop breeding, particularly to enhance adaptation to diverse environmental stresses. Variation in RSA and its plasticity influences above-ground growth by regulating water and nutrient acquisition, thereby supporting overall plant performance **[42–47]**. Temperature is another key environmental factor governing plant growth and development, and, in the absence of water limitation, often exerts a stronger influence than other environmental variables **[6]**.

Understanding how RSA responds to high-temperature stress requires evaluating genetically diverse wheat germplasm. The Multiple Synthetic Derivatives (MSD) population provides a valuable platform for this purpose, incorporating genetic diversity from *Aegilops tauschii* into a hexaploid wheat background **[48,34]**. In our previous study **[32]**, we developed a root phenotyping platform to analyze juvenile-stage RSA under contrasting temperature conditions and characterized the RSA of a representative MSD genotype, MSD417, and its recurrent parent, Norin 61 (N61). Although the MSD population shows substantial variation in various agronomic traits such as grain yield, biomass, heading time, stress tolerance, resource use efficiency, and grain quality **[34,49–52]**, their RSA characteristics remain poorly described. To address this gap, the present study expanded the analysis to include additional MSD genotypes (MSD034, MSD054, MSD296, and MSD392), as well as MSD417 and its breeding parent, N61. These MSD genotypes have been previously reported to exhibit heat tolerance **[40,41]**. Using time-course imaging of root development, we tracked dynamic changes in RSA under high-temperature conditions and clearly distinguished genotypic differences in root and shoot trait performance.

The ANOVA results indicate that both genotype and environment contributed to variation in root and shoot traits, with a particularly strong influence of high-temperature stress (Table 1). This dominant environmental effect suggests that RSA in wheat is highly plastic and responsive to high-temperature stress, consistent with previous reports **[9,53]**. At the same time, the significant genotypic effects observed across most traits highlight substantial genetic diversity within the MSD population. Importantly, the occurrence of significant genotype × environment (G × E) interactions for several key traits indicates that genotypes differ in their capacity to adjust RSA under high-temperature conditions. Such differential responses are particularly relevant for breeding, as they enable the identification of genotypes that maintain favorable root traits under high-temperature stress. In contrast, the absence of significant genotypic variation in traits such as rooting depth (RD) and primary root length (PRL) suggests that these vertical growth parameters are more constrained by environmental conditions than by genetic variation within this population. This may reflect a conserved developmental program for primary root elongation, whereas lateral expansion is more plastic and genotype-dependent under stress.

Under control conditions, MSD417 and MSD034 developed more extensive root systems, exhibiting higher values for key architectural traits such as total root length (TRL), root system width (RSW), convex hull area (CHA), and second pair seminal root length (SPSRL) (Figures 1, 3A, 4, 5). Although high-temperature stress reduced overall root growth and masked some genotypic differences, these two genotypes maintained relatively robust root systems compared to the others. In contrast, MSD392 showed pronounced sensitivity to RSW and CHA under high-temperature stress (Figure 1K, L), whereas MSD054 appeared less affected, likely reflecting its comparatively limited root development under control conditions.

A key observation of this study is that high-temperature stress tended to have a greater impact on traits related to lateral root system expansion, particularly SPSRL, RSW, and CHA, compared with vertical traits such as RD and PRL (Figures 1, 3, 4). These lateral root-related traits generally exhibited greater changes in stress index analyses (Figure 4), suggesting that horizontal root expansion may be an important component of RSA plasticity under high-temperature stress. This pattern is consistent with previous studies showing that lateral root development is responsive to environmental stresses and contributes to adaptive root plasticity **[44,54,16]**. Such lateral expansion may enhance the plant’s ability to exploit heterogeneous soil resources and maintain functional root systems under stress conditions **[47,55]**.

The stress index-based PCA further highlights these genotype-specific response strategies by integrating multiple performance and sensitivity metrics (Figure 5B). Such integrative approaches have been widely used to evaluate stress tolerance by combining multiple traits into composite indices **[56,57]**. Genotypes such as MSD417 and MSD034 were closely associated with performance-related indices (STI and MRP), indicating their ability to maintain functional root architecture under stress. In contrast, MSD392 was aligned with sensitivity-related indices (PR and SSPI), reflecting greater susceptibility to high-temperature stress.

Notably, MSD054 showed low values across both tolerance and sensitivity indices (Figures 4, 5B), suggesting a distinct response pattern characterized by limited responsiveness rather than strong tolerance or susceptibility. Its apparent stability may not reflect active stress tolerance but rather a limited baseline root system that undergoes minimal change under stress. This observation highlights an important distinction between true tolerance, defined by the maintenance of high performance under stress, and passive stability resulting from inherently low growth potential. Similar distinctions between tolerance and avoidance or low responsiveness have been discussed in stress physiology studies **[58,59]**. Such distinctions are crucial when selecting genotypes for breeding programs, as stable but low-performing genotypes may not contribute to yield improvement under stress conditions.

Taken together, the evaluated genotypes in this study can be broadly categorized into distinct response strategies: (i) high-performing and tolerant genotypes (MSD417 and MSD034), which maintain robust RSA under stress; (ii) stress-sensitive genotypes (e.g., MSD392), which exhibit pronounced reductions in root traits; and (iii) low-performing but relatively stable genotypes (e.g., MSD054), which show limited responsiveness to stress. This classification aligns with conceptual frameworks describing plant stress responses as combinations of tolerance, sensitivity, and avoidance strategies **[58,60]**. These results demonstrate that the MSD population encompasses a wide spectrum of adaptive strategies that can be effectively resolved through integrated phenotypic and index-based analyses.

From a breeding perspective, these findings emphasize the importance of selecting genotypes that combine high baseline performance with the capacity to maintain root system functionality under stress. Traits associated with lateral root expansion, such as SPSRL, RSW, and CHA, emerge as promising targets for selection. Previous studies have demonstrated that root architectural traits are closely linked to resource acquisition efficiency and yield stability under stress conditions **[61,21,62]**. Furthermore, integrating versatile, high-throughput root phenotyping platforms with genetically diverse populations, including MSD, provides a powerful framework for identifying loci associated with stress-adaptive RSA traits **[31,23,32]**. Further studies should focus on validating these traits under field conditions and linking them to yield stability to fully exploit their potential in wheat improvement.

## 4. Materials and Methods

### 4.1. Plant material

Seeds of bread wheat cultivar ‘Norin61’ (hereafter referred to as N61) were kindly provided by Prof. Hiroyuki Tanaka (Faculty of Agriculture, Tottori University, Tottori, Japan). Seeds of MSD genotypes (MSD034, MSD054, MSD296, MSD392, and MSD417) were kindly provided by Dr. Yasir Serag Alnor Gorafi (Arid Land Research Center, Tottori University, Tottori, Japan). The seeds were multiplied in pots at a glasshouse facility in the Faculty of Agriculture, Tottori University, Tottori, Japan, and fully mature seeds were harvested in 2024 and used for the experiments.

### 4.2. Plant growth

Wheat seedlings were grown using a calico cloth-based growth panel system described elsewhere **[32]**. Briefly, the growth panel consisted of an acrylic board (31 cm height, 30 cm width, and 0.5 cm thickness) layered with two sheets of paper towel (Kimtowel, Nippon Paper Crecia, Tokyo, Japan) soaked with nutrient solution (2000-fold diluted HYPONeX solution, Hyponex Japan, Osaka, Japan), and a black calico cloth. Germinated seeds were secured on the cloth with rubber bands 2 cm below the edge of the panel top. The stack was further covered with a high-density polyethylene sheet (0.05 mm width), followed by an additional layer of black calico cloth, and the assembly was secured with clips. The assembled panels were placed vertically in a plastic tray filled with a 2000-fold-diluted HYPONeX nutrient solution (to a depth of 3 cm). The submerged paper towel and calico cloth ensured a steady supply of moisture and nutrients to seedlings via capillary action. The unit was placed in a growth chamber (14/10 h light/dark regimes, 22/18 °C day/night temperatures, light intensity of approximately 350 µmol m−2 s−1, and relative humidity of 50/60 %). The panels were incubated at normal temperature conditions (14/10 h light/dark regimes, 22 °C/18 °C day/night temperatures), and after four days, half of the panels were transferred to a stress chamber at 42 °C/18 °C day/night temperatures) for another 4 days, while the control plants were maintained at normal temperatures (22 °C/18 °C).

### 4.3. Root image processing and biomass measurement

Root and shoot morphology was photographed daily using a bench-top photograph system (model ET16 Plus, CZUR Tech Co. Ltd., Oriental Science and Technology Building, No. 16, Keyuan Road, Nanashan District, Shenzen, China) as described previously **[32]**. On the eighth day, the root and shoot were harvested separately, weighed on an electronic balance to record fresh weight, and then dried for 3 days in a drying oven at 80 °C to record their dry weight. Root images were processed using ImageJ (v1.54g) integrated with the SmartRoot (v4.21) plugin **[63]** to quantify total and individual root lengths, along with the convex hull area. Additional traits, including rooting depth, root system width, seminal root angle, and shoot height, were measured directly in ImageJ.

### 4.4. Stress indices calculation

Representative stress indices for plant growth, percent reduction (PR) **[64]**, stress tolerance index (STI) **[65]**, stress susceptibility percentage index (SSPI) **[66]**, and mean relative performance (MRP) **[67]** ― were calculated in Microsoft Excel (Microsoft 365, Microsoft, Redmond, WA, USA) according to the following formulas:

PR = (Yc – Ys) / Yc × 100

STI = (Ys × Yc) / (Yc)^2^

SSPI = (Yc – Ys) / 2 × (Xc) × 100

MRP = (Ys / Xs) + (Yc / Xc)

where Yc and Ys represent the mean values of a given trait for a genotype under control and high-temperature conditions, respectively, and Xc and Xs represent the corresponding mean values across all genotypes under control and high-temperature conditions, respectively.

### 4.5. Statistical analysis

Preliminary data formatting was performed in Microsoft Excel (Microsoft 365). All subsequent statistical analyses and visualizations were conducted using the R statistical environment (v4.5.1; R Core Team, 2025). Tukey’s HSD method was conducted using the agricolae package (version 1.3.7), while Student’s t-test was performed using the R base package stats (version 4.5.1). Hierarchical clustering analysis (HCA), pairwise

phenotypic distance analysis (Euclidean distances), and principal component analysis (PCA) were performed by pheatmap (version 1.0.13), R base package stats (version 4.5.1), and FactoMineR (version 2.12) packages, respectively. All custom R scripts developed for this study are available in Supplementary Document S1.

## 5. Conclusions

Based on a set of representative genotypes, the MSD wheat population provides a powerful platform for studying root system architecture (RSA) under high-temperature stress. Substantial plasticity in the RSA was observed within this genetically diverse population, with lateral root expansion traits responding more than vertical growth traits. By integrating phenotypic analyses and stress indices, we identified distinct genotypic response strategies. In particular, MSD417 and MSD034 maintained higher root performance under stress; MSD392 was sensitive; and MSD054 responded poorly. These findings highlight the importance of distinguishing true stress tolerance from passive stability and emphasize the value of lateral root traits as targets for selection. Combining germplasm, root phenotyping, and analysis frameworks provides a basis for studying RSA-mediated stress adaptations. Future research should validate these findings under field conditions and link RSA traits to yield stability.

## Supporting information

Supplementary Document R script

## Supplementary Materials

The following supporting information is available: Supplementary file S1: R-scripts for the processing of data for root system architecture, Supplementary file S2: Supplementary Tables S1-S6.

## Author Contributions

Conceptualization, S.M.M.I. and K.A.; methodology, S.M.M.I. and KA; software, S.M.M.I.; validation, S.M.M.I., I.S.A.T., and K.A.; formal analysis, S.M.M.I.; investigation, S.M.M.I.; resources, S.M.M.I.; data curation, S.M.M.I.; writing—original draft preparation, S.M.M.I.; writing—review and editing, I.S.A.T. and K.A.; visualization, S.M.M.I. and K.A.; supervision, K.A.; project administration, K.A.; funding acquisition, K.A. All authors have read and agreed to the published version of the manuscript.

## Funding

This research was funded by the Project Marginal Region Agriculture, the Arid Land Research Center, Tottori University, and the IPDRE Program, Tottori University.

## Data Availability Statement

The root images of wheat N61, MSD034, MSD054, MSD296, MSD392 and MSD417 are deposited in the Zenodo data repository under https://doi.org/10.5281/zenodo.19708100, https://doi.org/10.5281/zenodo.19708508, https://doi.org/10.5281/zenodo.19708809, https://doi.org/10.5281/zenodo.19709019, https://doi.org/10.5281/zenodo.19709196, and https://doi.org/10.5281/zenodo.19709468, respectively. The other original contributions presented in the study are included in the article/supplementary material.

## Acknowledgments

We are grateful to Prof Hiroyuki Tanaka (Faculty of Agriculture, Tottori University, Tottori, Japan) and Dr Yasir Serag Alnor Gorafi (Graduate School of Agriculture, Kyoto University, Kyoto, Japan) for providing seeds of wheat cultivar Norin 61 and the MSD genotype MSD417, respectively. We thank the staff members of the Laboratory of Molecular and Cellular Biology, Faculty of Agriculture, Tottori University, for their technical support in the laboratory.

## Conflicts of Interest

The authors declare no conflict of interest.

## Notes

### Competing Interest Statement

The authors have declared no competing interest.

