## Supplementary Tables for "Root system architecture responses to high-temperature stress in synthetic-derived wheat lines reveal distinct adaptive patterns"

Supplementary Table S1. Descriptive statistics of root and shoot traits of the tested genotypes at the contrasting temperatures.

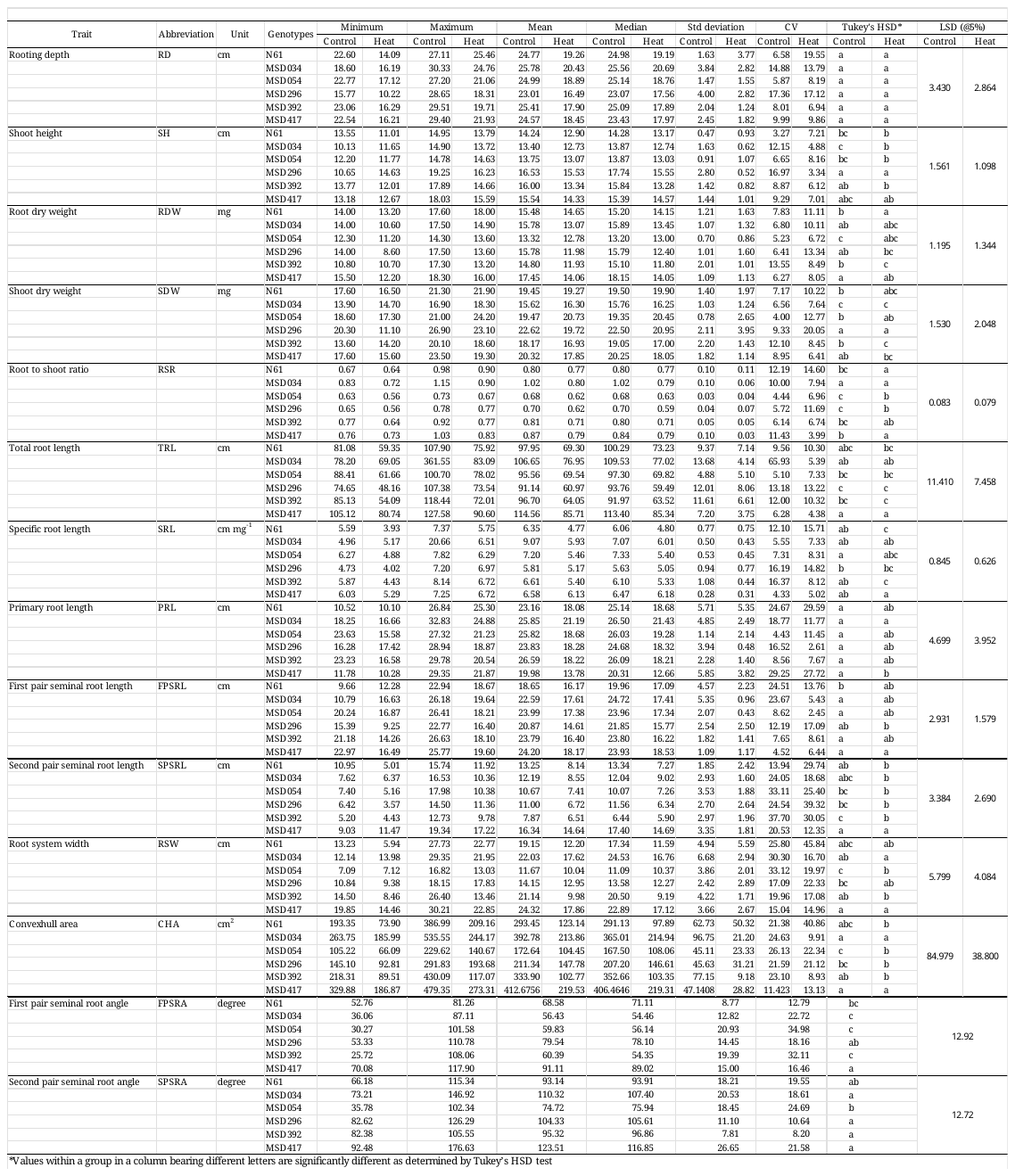

Supplementary Table S2. Percent reduction of different root and shoot traits under high temperature conditions compared to control conditions in wheat genotype.

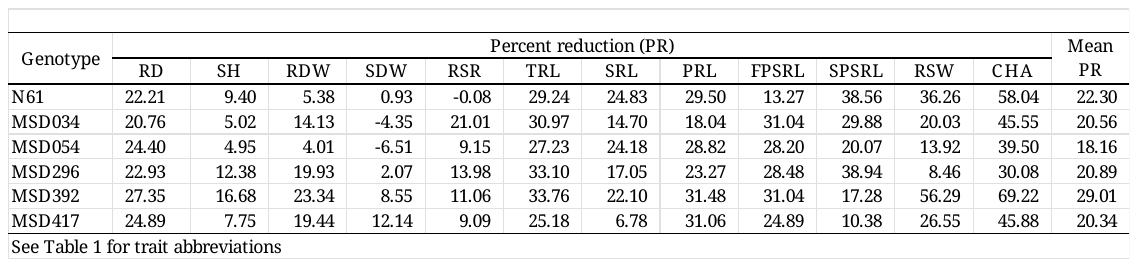

Supplementary Table S3. Stress susceptibility percentage index of different root and shoot traits under high temperature conditions compared to control conditions in wheat genotype

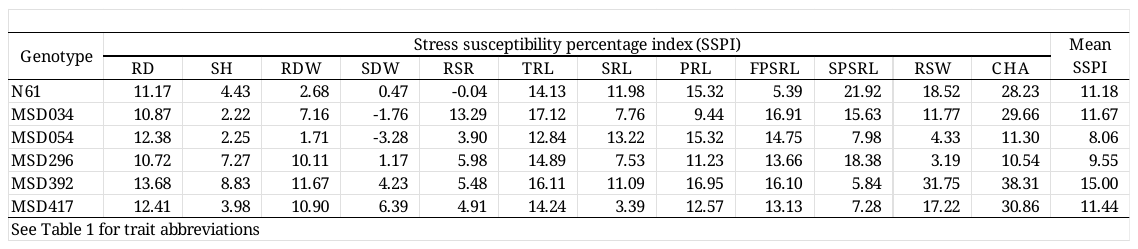

Supplementary Table S4. Mean relative performance of different root and shoot traits under high temperature condition compared to control conditions in wheat genotype

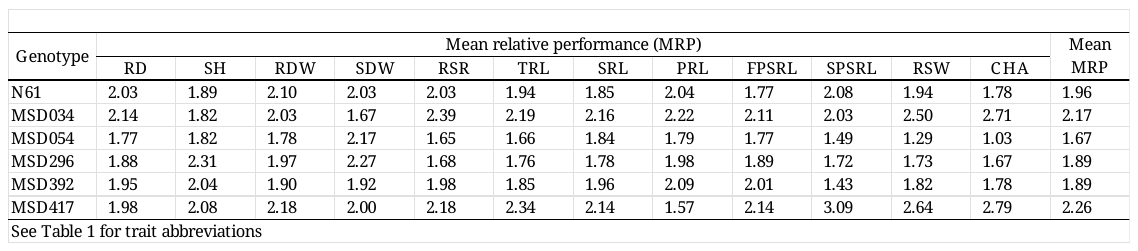

Supplementary Table S5. Stress tolerance index of different root and shoot traits under high temperature condition compared to control conditions in wheat genotype.

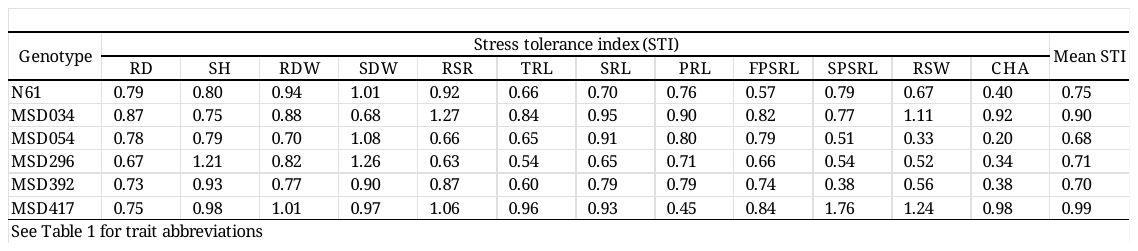
