## Supplementary Document R script for "Root system architecture responses to high-temperature stress in synthetic-derived wheat lines reveal distinct adaptive patterns"

R-scripts for the processing of data for root system architecture

**List of scripts:**

Script number 1. Two-way analysis of variance (ANOVA) for different root and shoot traits at day eight in contrasting temperature conditions. (Page: 01-03).

Script number 2. Tukeys HSD post hoc test for different root and shoot traits of different genotypes at day eight in contrasting temperature conditions. (Page: 03-06).

Script number 3. Student’s t-test for different root and shoot traits of different genotypes in contrasting temperature conditions. (Page: 06-09).

Script number 4. Tukeys HSD post hoc test for first pair seminal root angle (FPSRA) and second pair seminal root angle (SPSRA) of different genotypes at day eight (Page: 09-10).

Script number 5. Hierarchical cluster analysis, double dendrogram and heatmap of wheat genotypes on measured root and shoot traits (Page: 10-11).

Script number 6. Principal Component Analysis (PCA) biplot of wheat genotypes under contrasting temperature conditions (Page: 11-12).

Script number 7. Principal Component Analysis (PCA) biplot of wheat genotypes for stress indices (Page: 12-13).

Script number 8. Tukeys HSD post hoc test for different root and shoot traits of different genotypes at day eight in control condition. (Page: 13-15).

Script number 9. Tukeys HSD post hoc test for different root and shoot traits of different genotypes at day eight in high temperature stress condition. (Page: 15-18).

Script number 1.

Two-way analysis of variance (ANOVA) for different root and shoot traits at day eight in contrasting temperature conditions.

\# Install and load necessary packages

library(agricolae)

library(dplyr)

library(ggplot2)

library(broom)

library(tidyverse)

\# Memory cleaning and folder preparation

\# Clean up the system brain if any

rm(list=ls())

\# Select the .csv file

setwd("D:/251006_R_analysis")

FinalDayData <-read.csv("D:/251006_R_analysis/251230_FinalDayData_all_Genotype.csv")

Variables <- c("RD", "SH", "RDW", "SDW", "RSR","TRL", "SRL", "PRL", "FPSRL", "SPSRL",

"RW")

\# Select the categorical variables

FinalDayData$Genotype <- as.factor(FinalDayData$Genotype)

FinalDayData$Condition <- as.factor(FinalDayData$Condition)

\## ANOVA for Rooting depth (RD)

ANOVA_RD <- aov(RD ~ Genotype * Condition, data = FinalDayData)

\# Data framing for downloading ANOVA result

ANOVA_summary_RD <- tidy(ANOVA_RD)

\# Save data as CSV file

write.csv(ANOVA_summary_RD, "RD_ANOVA_result.csv", row.names = FALSE)

\## ANOVA for Shoot height (SH)

ANOVA_SH <- aov(SH ~ Genotype * Condition, data = FinalDayData)

\# Data framing for downloading ANOVA result

ANOVA_summary_SH <- tidy(ANOVA_SH)

\# Save data as CSV file

write.csv(ANOVA_summary_SH, "SH_ANOVA_result.csv", row.names = FALSE)

\## ANOVA for Root dry weight (RDW)

ANOVA_RDW <- aov(RDW ~ Genotype * Condition, data = FinalDayData)

\# Data framing for downloading ANOVA result

ANOVA_summary_RDW <- tidy(ANOVA_RDW)

\# Save data as CSV file

write.csv(ANOVA_summary_RDW, "RDW_ANOVA_result.csv", row.names = FALSE)

\## ANOVA for Shoot dry weight (SDW)

ANOVA_SDW <- aov(SDW ~ Genotype * Condition, data = FinalDayData)

\# Data framing for downloading ANOVA result

ANOVA_summary_SDW <- tidy(ANOVA_SDW)

\# Save data as CSV file

write.csv(ANOVA_summary_SDW, "SDW_ANOVA_result.csv", row.names = FALSE)

\## ANOVA for Root to shoot ratio (RSR)

ANOVA_RSR <- aov(RSR ~ Genotype * Condition, data = FinalDayData)

\# Data framing for downloading ANOVA result

ANOVA_summary_RSR <- tidy(ANOVA_RSR)

\# Save data as CSV file

write.csv(ANOVA_summary_RSR, "RSR_ANOVA_result.csv", row.names = FALSE)

\## ANOVA for total root length (TRL)

ANOVA_TRL <- aov(TRL ~ Genotype * Condition, data = FinalDayData)

\# Data framing for downloading ANOVA result

ANOVA_summary_TRL <- tidy(ANOVA_TRL)

\# Save data as CSV file

write.csv(ANOVA_summary_TRL, "TRL_ANOVA_result.csv", row.names = FALSE)

\## ANOVA for Specific root length (SRL)

ANOVA_SRL <- aov(SRL ~ Genotype * Condition, data = FinalDayData)

\# Data framing for downloading ANOVA result

ANOVA_summary_SRL <- tidy(ANOVA_SRL)

\# Save data as CSV file

write.csv(ANOVA_summary_SRL, "SRL_ANOVA_result.csv", row.names = FALSE)

\## ANOVA for Primary root length (PRL)

ANOVA_PRL <- aov(PRL ~ Genotype * Condition, data = FinalDayData)

\# Data framing for downloading ANOVA result

ANOVA_summary_PRL <- tidy(ANOVA_PRL)

\# Save data as CSV file

write.csv(ANOVA_summary_PRL, "PRL_ANOVA_result.csv", row.names = FALSE)

\## ANOVA for First pair seminal root length (FPSRL)

ANOVA_FPSRL <- aov(FPSRL ~ Genotype * Condition, data = FinalDayData)

\# Data framing for downloading ANOVA result

ANOVA_summary_FPSRL <- tidy(ANOVA_FPSRL)

\# Save data as CSV file

write.csv(ANOVA_summary_FPSRL, "FPSRL_ANOVA_result.csv", row.names = FALSE)

\## ANOVA for Second pair seminal root length (SPSRL)

ANOVA_SPSRL <- aov(SPSRL ~ Genotype * Condition, data = FinalDayData)

\# Data framing for downloading ANOVA result

ANOVA_summary_SPSRL <- tidy(ANOVA_SPSRL)

\# Save data as CSV file

write.csv(ANOVA_summary_SPSRL, "SPSRL_ANOVA_result.csv", row.names = FALSE)

\## ANOVA: root system width (RSW)

ANOVA_RW <- aov(RW ~ Genotype * Condition, data = FinalDayData)

\# Data framing for downloading ANOVA result

ANOVA_summary_RW <- tidy(ANOVA_RW)

\# Save data as CSV file

write.csv(ANOVA_summary_RW, "RW_ANOVA_result.csv", row.names = FALSE)

\## ANOVA: convexhull area (CHA)

ANOVA_CHA <- aov(CHA ~ Genotype * Condition, data = FinalDayData)

\# Data framing for downloading ANOVA result

ANOVA_summary_CHA <- tidy(ANOVA_CHA)

\# Save data as CSV file

write.csv(ANOVA_summary_CHA, "CHA_ANOVA_result.csv", row.names = FALSE)

Script number 2.

Tukeys HSD post hoc test for different root and shoot traits of different genotypes at day eight in contrasting temperature conditions.

\# Install and load necessary packages

library(agricolae)

library(dplyr)

library(ggplot2)

library(broom)

library(tidyverse)

\# Memory cleaning and folder preparation

\# (1) Clean up the system brain if any

rm(list=ls())

\# (2) Select the .csv file

setwd("D:/251006_R_analysis")

FinalDayData <-read.csv("D:/251006_R_analysis/251230_FinalDayData_all_Genotype.csv

Variables <- c("RD", "SH", "RDW", "SDW", "RSR","TRL", "SRL", "PRL", "FPSRL", "SPSRL",

"RW", "CHA")

\## ANOVA: Rooting depth (RD)

ANOVA_RD <- aov(RD ~ Genotype, data = FinalDayData)

\## Tukey's HSD post-hoc test: rooting depth (RD)

HSD_result_RD <- HSD.test(ANOVA_RD, trt = "Genotype", alpha = 0.05, console = TRUE)

\# Data framing to download

HSD_group_RD <- HSD_result_RD$groups

\# Save the data frame as CSV file

write.csv(HSD_group_RD, file = "RD_HSD_result.csv")

\## ANOVA: shoot height (SH)

ANOVA_SH <- aov(SH ~ Genotype, data = FinalDayData)

\## Tukey's HSD post-hoc test: shoot height (SH)

HSD_result_SH <- HSD.test(ANOVA_SH, trt = "Genotype", alpha = 0.05, console = TRUE)

\# Data framing to download

HSD_group_SH <- HSD_result_SH$groups

\# Save the data frame as CSV file

write.csv(HSD_group_SH, file = "SH_HSD_result.csv")

\## ANOVA: root dry weight (RDW)

ANOVA_RDW <- aov(RDW ~ Genotype, data = FinalDayData)

\## Tukey's HSD post-hoc test: root dry weight (RDW)

HSD_result_RDW <- HSD.test(ANOVA_RDW, trt = "Genotype", alpha = 0.05, console = TRUE)

\# Data framing to download

HSD_group_RDW <- HSD_result_RDW$groups

\# Save the data frame as CSV file

write.csv(HSD_group_RDW, file = "RDW_HSD_result.csv")

\## ANOVA: shoot dry weight (SDW)

ANOVA_SDW <- aov(SDW ~ Genotype, data = FinalDayData)

\## Tukey's HSD post-hoc test: shoot dry weight (SDW)

HSD_result_SDW <- HSD.test(ANOVA_SDW, trt = "Genotype", alpha = 0.05, console = TRUE)

\# Data framing to download

HSD_group_SDW <- HSD_result_SDW$groups

\# Save the data frame as CSV file

write.csv(HSD_group_SDW, file = "SDW_HSD_result.csv")

\## ANOVA: root to shoot ratio (RSR)

ANOVA_RSR <- aov(RSR ~ Genotype, data = FinalDayData)

\## Tukey's HSD post-hoc test: root to shoot ratio (RSR)

HSD_result_RSR <- HSD.test(ANOVA_RSR, trt = "Genotype", alpha = 0.05, console = TRUE)

\# Data framing to download

HSD_group_RSR <- HSD_result_RSR$groups

\# Save the data frame as CSV file

write.csv(HSD_group_RSR, file = "RSR_HSD_result.csv")

\## ANOVA: total root length (TRL)

ANOVA_TRL <- aov(TRL ~ Genotype, data = FinalDayData)

\## Tukey's HSD post-hoc test: total root length (TRL)

HSD_result_TRL <- HSD.test(ANOVA_TRL, trt = "Genotype", alpha = 0.05, console = TRUE)

\# Data framing to download

HSD_group_TRL <- HSD_result_TRL$groups

\# Save the data frame as CSV file

write.csv(HSD_group_TRL, file = "TRL_HSD_result.csv")

\## ANOVA: specific root length (SRL)

ANOVA_SRL <- aov(SRL ~ Genotype, data = FinalDayData)

\## Tukey's HSD post-hoc test: specific root length (SRL)

HSD_result_SRL <- HSD.test(ANOVA_SRL, trt = "Genotype", alpha = 0.05, console = TRUE)

\# Data framing to download

HSD_group_SRL <- HSD_result_SRL$groups

\# Save the data frame as CSV file

write.csv(HSD_group_SRL, file = "SRL_HSD_result.csv")

\## ANOVA: primary root length (PRL)

ANOVA_PRL <- aov(PRL ~ Genotype, data = FinalDayData)

\## Tukey's HSD post-hoc test: primary root length (PRL)

HSD_result_PRL <- HSD.test(ANOVA_PRL, trt = "Genotype", alpha = 0.05, console = TRUE)

\# Data framing to download

HSD_group_PRL <- HSD_result_PRL$groups

\# Save the data frame as CSV file

write.csv(HSD_group_PRL, file = "PRL_HSD_result.csv")

\## ANOVA: first pair seminal root length (FPSRL)

ANOVA_FPSRL <- aov(FPSRL ~ Genotype, data = FinalDayData)

\## Tukey's HSD post-hoc test: first pair seminal root length (FPSRL)

HSD_result_FPSRL <- HSD.test(ANOVA_FPSRL, trt = "Genotype", alpha = 0.05, console = TRUE)

\# Data framing to download

HSD_group_FPSRL <- HSD_result_FPSRL$groups

\# Save the data frame as CSV file

write.csv(HSD_group_FPSRL, file = "FPSRL_HSD_result.csv")

\## ANOVA: second pair seminal root length (SPSRL)

ANOVA_SPSRL <- aov(SPSRL ~ Genotype, data = FinalDayData)

\## Tukey's HSD post-hoc test: second pair seminal root length (SPSRL)

HSD_result_SPSRL <- HSD.test(ANOVA_SPSRL, trt = "Genotype", alpha = 0.05, console = TRUE)

\# Data framing to download

HSD_group_SPSRL <- HSD_result_SPSRL$groups

\# Save the data frame as CSV file

write.csv(HSD_group_SPSRL, file = "SPSRL_HSD_result.csv")

\## ANOVA: root system width (RSW)

ANOVA_RW <- aov(RW ~ Genotype, data = FinalDayData)

\## Tukey's HSD post-hoc test: root system width (RSW)

HSD_result_RW <- HSD.test(ANOVA_RW, trt = "Genotype", alpha = 0.05, console = TRUE)

\# Data framing to download

HSD_group_RW <- HSD_result_RW$groups

\# Save the data frame as CSV file

write.csv(HSD_group_RW, file = "RW_HSD_result.csv")

\## ANOVA: convexhull area (CHA)

ANOVA_CHA <- aov(CHA ~ Genotype, data = FinalDayData)

\## Tukey's HSD post-hoc test: convexhull area (CHA)

HSD_result_CHA <- HSD.test(ANOVA_CHA, trt = "Genotype", alpha = 0.05, console = TRUE)

\# Data framing to download

HSD_group_CHA <- HSD_result_CHA$groups

\# Save the data frame as CSV file

write.csv(HSD_group_CHA, file = "CHA_HSD_result.csv")

Script number 3.

Student’s *t*-test for different root and shoot traits of different genotypes in contrasting temperature conditions**.**

\## Installed and load the necessary packages

library(agricolae)

library(dplyr)

library(ggplot2)

library(broom)

library(tidyverse)

\## Memory cleaning and folder preparation

\# Clean up the system brain if any

rm(list=ls())

\# Select the .csv file

setwd("D:/251006_R_analysis")

FinalDayData<-read.csv("D:/251006_R_analysis/251204_FinalDayData_all_Genotype.csv")

\# t-test of N61_C vs. N61_H

\# create a subset data of N61 in two conditions

subset_N61 <- subset(FinalDayData, Genotype %in% c("N61_C", "N61_H"))

Variables <- c("RD", "SH", "RDW", "SDW", "RSR","TRL", "SRL", "PRL", "FPSRL", "SPSRL",

"RSW", "CHA")

\## Run t-Test

t_Test_N61 <- subset_N61 %>%

pivot_longer(cols = all_of(Variables),

names_to = "Variable",

values_to = "Value") %>%

group_by(Variable) %>%

summarize(broom::tidy(t.test(Value ~ Genotype))) %>%

ungroup() %>%

rename(

t_statistic = statistic,

p_value = p.value,

degrees_of_freedom = parameter,

conf_interval_low = conf.low,

conf_interval_high = conf.high,

mean_group1 = estimate1,

mean_group2 = estimate2)

\# View the final, combined data frame

print(t_Test_N61)

\# Save the results to a CSV

write.csv(t_Test_N61, "t_Test_N61_results.csv", row.names = FALSE)

\## t-test of MSD034_C vs. MSD034_H

subset_MSD034 <- subset(FinalDayData, Genotype %in% c("MSD034_C", "MSD034_H"))

\# Run t-Test

t_Test_MSD034 <- subset_MSD034 %>%

pivot_longer(

cols = all_of(Variables),

names_to = "Variable",

values_to = "Value") %>%

group_by(Variable) %>%

summarize(broom::tidy(t.test(Value ~ Genotype))) %>%

ungroup() %>%

rename(t_statistic = statistic,

p_value = p.value,

degrees_of_freedom = parameter,

conf_interval_low = conf.low,

conf_interval_high = conf.high,

mean_group1 = estimate1,

mean_group2 = estimate2)

\# Save the results as CSV

write.csv(t_Test_MSD034, "t_Test_MSD034_results.csv", row.names = FALSE)

\## t-test of MSD054_C vs. MSD054_H

subset_MSD054 <- subset(FinalDayData, Genotype %in% c("MSD054_C", "MSD054_H"))

\# Run t-Test

t_Test_MSD054 <- subset_MSD054 %>%

pivot_longer(

cols = all_of(Variables),

names_to = "Variable",

values_to = "Value") %>%

group_by(Variable) %>%

summarize(broom::tidy(t.test(Value ~ Genotype))) %>%

ungroup() %>%

rename(t_statistic = statistic,

p_value = p.value,

degrees_of_freedom = parameter,

conf_interval_low = conf.low,

conf_interval_high = conf.high,

mean_group1 = estimate1,

mean_group2 = estimate2)

\# Save the results as CSV

write.csv(t_Test_MSD054, "t_Test_MSD054_results.csv", row.names = FALSE)

\## t-test of MSD296_C vs. MSD296_H

subset_MSD296 <- subset(FinalDayData, Genotype %in% c("MSD296_C", "MSD296_H"))

\# Run t-Test

t_Test_MSD296 <- subset_MSD296 %>%

pivot_longer(

cols = all_of(Variables),

names_to = "Variable",

values_to = "Value") %>%

group_by(Variable) %>%

summarize(broom::tidy(t.test(Value ~ Genotype))) %>%

ungroup() %>%

rename(t_statistic = statistic,

p_value = p.value,

degrees_of_freedom = parameter,

conf_interval_low = conf.low,

conf_interval_high = conf.high,

mean_group1 = estimate1,

mean_group2 = estimate2)

\# Save the results as CSV

write.csv(t_Test_MSD296, "t_Test_MSD296_results.csv", row.names = FALSE)

\## t-test of MSD392_C vs. MSD392_H

subset_MSD392 <- subset(FinalDayData, Genotype %in% c("MSD392_C", "MSD392_H"))

\# Run t-Test

t_Test_MSD392 <- subset_MSD392 %>%

pivot_longer(

cols = all_of(Variables),

names_to = "Variable",

values_to = "Value") %>%

group_by(Variable) %>%

summarize(broom::tidy(t.test(Value ~ Genotype))) %>%

ungroup() %>%

rename(t_statistic = statistic,

p_value = p.value,

degrees_of_freedom = parameter,

conf_interval_low = conf.low,

conf_interval_high = conf.high,

mean_group1 = estimate1,

mean_group2 = estimate2)

\# Save the results as CSV

write.csv(t_Test_MSD392, "t_Test_MSD392_results.csv", row.names = FALSE)

\## t-test of MSD417_C vs. MSD417_H

subset_MSD417 <- subset(FinalDayData, Genotype %in% c("MSD417_C", "MSD417_H"))

\# Run t-Test

t_Test_MSD417 <- subset_MSD417 %>%

pivot_longer(

cols = all_of(Variables),

names_to = "Variable",

values_to = "Value") %>%

group_by(Variable) %>%

summarize(broom::tidy(t.test(Value ~ Genotype))) %>%

ungroup() %>%

rename(t_statistic = statistic,

p_value = p.value,

degrees_of_freedom = parameter,

conf_interval_low = conf.low,

conf_interval_high = conf.high,

mean_group1 = estimate1,

mean_group2 = estimate2)

\# Save the results as CSV

write.csv(t_Test_MSD417, "t_Test_MSD417_results.csv", row.names = FALSE)

Script number 4.

Tukeys HSD post hoc test for first pair seminal root angle (FPSRA) and second pair seminal root angle (SPSRA) of different genotypes at day eight

\## Install and load the necessary packages

library(agricolae)

library(dplyr)

library(ggplot2)

library(broom)

library(tidyverse)

\# Memory cleaning and folder preparation

\# Clean up the system brain if any

rm(list=ls())

\# Select .csv file

setwd("D:/251006_R_analysis")

FinalDayData <-read.csv("D:/251006_R_analysis/251210_SRA_5_Genotypes.csv")

Variables <- c("FPSRA", "SPSRA")

\# Tukeys HSD test for all genotypes

HSD_groups_FDD <-FinalDayData %>%

pivot_longer(cols = all_of(Variables),

names_to = "Variable",

values_to = "Value") %>%

group_by(Variable) %>%

nest() %>% mutate(model = map(data, ~ aov(Value ~ Genotype, data = .)),

hsd_result = map(model, ~ HSD.test(., trt = "Genotype", alpha = 0.05, console = FALSE)),

groups_df = map(hsd_result, ~ as.data.frame(.$groups) %>%

rownames_to_column(var = "Genotype"))) %>%

select(Variable, groups_df) %>%

unnest(cols = groups_df) %>%

rename(Mean = Value)

\# Save the results as CSV

write.csv(HSD_groups_FDD, "Tukeys_HSD_groups_FDD.csv", row.names = FALSE)

Script number 5.

Hierarchical cluster analysis, double dendrogram and heatmap of wheat genotypes on measured root and shoot traits

\# Install and load the necessary packages

library(dplyr)

library(pheatmap)

library(tibble)

\## Memory cleaning and folder preparation

\# Clean up the system brain if any

rm(list=ls())

\# Select the .csv file

setwd("D:/251006_R_analysis")

FinalDayData <-read.csv("D:/251006_R_analysis/260113_FinalDayData_all_Genotype.csv")

\# Calculate the mean of each treatment for each variable

Mean_data <- FinalDayData %>%

group_by(Genotype) %>%

summarise(across(RD:SPSRA, mean, .names = "{.col}")) %>%

as.data.frame()

\# Format the row name as Genotype

data_for_plot <- Mean_data %>% tibble::column_to_rownames("Genotype")

\# Prepare Plot A: Hierarchical Cluster Heatmap

Heatmap_A <- pheatmap(data_for_plot,

main = "Plot A: Hierarchical Cluster Heatmap",

scale = "column",

cluster_rows = TRUE, # Cluster the genotypes

cluster_cols = TRUE, # Cluster the variables

cutree_rows = 5, # This will cut the row dendrogram into 3 clusters, adjust if youneed more cluster

fontsize_row = 18,

fontsize_col = 18,

fontsize = 16,

fontsize_number = 14)

\# Save Plot as PNG

HeatMap_A <- paste0(format(Sys.Date(), "%y%m%d"),"_Hierarchical_Cluster_Heatmap_A", ".png")

Location_path <- file.path("D:/251006_R_analysis", HeatMap_A)

png(Location_path, width = 800, height = 1000, res = 100)

dev.off()

\## Preparation of Plot B: XY Plot

\# Scale the data by column

data_scaled <- scale(data_for_plot)

\# Calculate the Euclidean distance matrix between the rows (Genotype)

dist_matrix <- dist(data_scaled, method = "euclidean")

dist_matrix_full <- as.matrix(dist_matrix)

\# Get the ordering of the rows from Plot A's clustering

row_order <- Heatmap_A$tree_row$order

\# Re-order the distance matrix to match the clustering from Plot A

dist_matrix_ordered <- dist_matrix_full[row_order, row_order]

\# Plot the distance matrix

Heatmap_B <- pheatmap(dist_matrix_ordered,

main = "Plot B: Dissimilarity Matrix (XY Plot)",

cluster_rows = FALSE, cluster_cols = FALSE, fontsize_row = 18, fontsize_col = 18,

fontsize = 16, fontsize_number = 14)

\# Save Plot as PNG

HeatMap_B <- paste0(format(Sys.Date(), "%y%m%d"),"_Hierarchical_Cluster_Heatmap_B", ".png")

Location_path <- file.path("D:/251006_R_analysis", HeatMap_B)

png(Location_path, width = 620, height = 956, res = 100)

dev.off()

Script number 6.

Principal Component Analysis (PCA) biplot of wheat genotypes under contrasting temperature conditions

\## Install and download the necessary packages

library(FactoMineR)

library(factoextra)

library(dplyr)

library(ggpubr)

\# Memory cleaning and folder preparation

\# Clean up the system brain if any

rm(list=ls())

\# Select the .csv file

setwd("D:/251006_R_analysis")

FinalDayData <-read.csv("D:/251006_R_analysis/260113_FinalDayData_all_Genotype.csv")

\# Aggregate Data (Calculate Means) ---

FinalDayData_mean <- FinalDayData %>%

group_by(Genotype) %>%

summarise(across(where(is.numeric), mean)) %>%

select(-Replication)

\# Set Genotypes as row names

df_pca <- FinalDayData_mean %>% tibble::column_to_rownames("Genotype")

\## Run PCA analysis

PCA_Mean_FDD <- PCA(df_pca, graph = FALSE)

\# Ensure conditions is a factor

conditions <- as.factor(conditions)

\# Create the Biplot

PCA_Biplot <- fviz_pca_biplot(PCA_Mean_FDD,

geom.ind = c("point", "text"),

habillage = conditions, pointsize = 3,

col.var = "black", select.var = list(contrib = 10),

mean.point = FALSE, show.legend = FALSE,

repel = TRUE,

title = "PCA - Biplot: Control vs. Heat",

labelsize = 4) +

scale_color_manual(values = c("Control" = "green", "Heat" = "red")) +

scale_shape_manual(values = c("Control" = 16, "Heat" = 17)) +

theme(axis.title = element_text(size = 14),

axis.text = element_text(size = 14),

legend.text = element_text(size = 12),

legend.title = element_text(size = 14),

legend.position = "none")

\# Save the PCA plot as png

PCA_Bi_plot <- paste0(format(Sys.Date(), "%y%m%d"),"_PCA_Biplot_with_title", ".png")

Location_path <- file.path("D:/251006_R_analysis", PCA_Bi_plot)

ggsave(filename = Location_path,

plot = PCA_Biplot,

width = 2000,

height = 1200,

units = "px",

dpi = 300)

Script number 7.

Principal Component Analysis (PCA) biplot of wheat genotypes for stress indices

\## Install and download the necessary packages

library(FactoMineR)

library(factoextra)

library(dplyr)

library(ggpubr)

\# Memory cleaning and folder preparation

\# Clean up the system brain if any

rm(list=ls())

\# Select the .csv file

setwd("D:/251006_R_analysis")

STI <-read.csv("D:/251006_R_analysis/260411_STI.csv")

\# Aggregate Data (Calculate Means) ---

FinalDayData_mean <- STI %>%

group_by(Genotype) %>%

summarise(across(where(is.numeric), mean)) %>%

select(-Replication)

\# Set Genotypes as row names

df_pca <- STI %>% tibble::column_to_rownames("Genotype")

\## Run PCA analysis

PCA_ STI <- PCA(df_pca, graph = FALSE)

\# Ensure conditions is a factor

conditions <- as.factor(conditions)

\# Create the Biplot

PCA_Biplot <- fviz_pca_biplot(PCA_ STI,

geom.ind = c("point", "text"),

habillage = conditions, pointsize = 3,

col.var = "black", select.var = list(contrib = 10),

mean.point = FALSE, show.legend = FALSE,

repel = TRUE,

title = "PCA_STI",

labelsize = 4) +

theme(axis.title = element_text(size = 14),

axis.text = element_text(size = 14),

legend.text = element_text(size = 12),

legend.title = element_text(size = 14),

legend.position = "none")

\# Save the PCA plot as png

PCA_Bi_plot <- paste0(format(Sys.Date(), "%y%m%d"),"_PCA_Biplot_with_title", ".png")

Location_path <- file.path("D:/251006_R_analysis", PCA_Bi_plot)

ggsave(filename = Location_path,

plot = PCA_Biplot,

width = 2000,

height = 1200,

units = "px",

dpi = 300)

Script number 8.

Tukeys HSD post hoc test for different root and shoot traits of different genotypes at day eight in control condition.

\## Install and load the necessary packageslibrary(agricolae)

library(dplyr)

library(ggplot2)

library(broom)

library(tidyverse)

\# Memory cleaning and folder preparation

\# Clean up the system brain if any

rm(list=ls())

\# Select csv file by the following command

setwd("D:/251006_R_analysis")

FinalDayData_all <-read.csv("D:/251006_R_analysis/251230_FinalDayData_all_Genotype.csv")

\# Create a sub-set data for Control condition

FinalDayData <- subset(FinalDayData_all, Genotype %in% c("N61_C", "MSD034_C", "MSD054_C", "MSD296_C", "MSD392_C", "MSD417_C"))

Variables <- c("RD", "SH", "RDW", "SDW", "RSR","TRL", "SRL", "PRL", "FPSRL", "SPSRL",

"RW", "CHA", "FPSRA", "SPSRA")

\## ANOVA: rooting depth (RD)

ANOVA_RD <- aov(RD ~ Genotype, data = FinalDayData)

\## Tukey's HSD post-hoc test: rooting depth (RD)

HSD_result_RD <- HSD.test(ANOVA_RD, trt = "Genotype", alpha = 0.05, console = TRUE)

\# Data framing to download

HSD_group_RD <- HSD_result_RD$groups

\# Save the data frame as CSV file

write.csv(HSD_group_RD, file = "RD_HSD_result.csv")

\## ANOVA: shoot height (SH)

ANOVA_SH <- aov(SH ~ Genotype, data = FinalDayData)

\## Tukey's HSD post-hoc test: shoot height (SH)

HSD_result_SH <- HSD.test(ANOVA_SH, trt = "Genotype", alpha = 0.05, console = TRUE)

\# Data framing to download

HSD_group_SH <- HSD_result_SH$groups

\# Save the data frame as CSV file

write.csv(HSD_group_SH, file = "SH_HSD_result.csv")

\## ANOVA: root dry weight (RDW)

ANOVA_RDW <- aov(RDW ~ Genotype, data = FinalDayData)

\## Tukey's HSD post-hoc test: root dry weight (RDW)

HSD_result_RDW <- HSD.test(ANOVA_RDW, trt = "Genotype", alpha = 0.05, console = TRUE)

\# Data framing to download

HSD_group_RDW <- HSD_result_RDW$groups

\# Save the data frame as CSV file

write.csv(HSD_group_RDW, file = "RDW_HSD_result.csv")

\## ANOVA: shoot dry weight (SDW)

ANOVA_SDW <- aov(SDW ~ Genotype, data = FinalDayData)

\## Tukey's HSD post-hoc test: shoot dry weight (SDW)

HSD_result_SDW <- HSD.test(ANOVA_SDW, trt = "Genotype", alpha = 0.05, console = TRUE)

\# Data framing to download

HSD_group_SDW <- HSD_result_SDW$groups

\# Save the data frame as CSV file

write.csv(HSD_group_SDW, file = "SDW_HSD_result.csv")

\## ANOVA: root to shoot ratio (RSR)

ANOVA_RSR <- aov(RSR ~ Genotype, data = FinalDayData)

\## Tukey's HSD post-hoc test: root to shoot ratio (RSR)

HSD_result_RSR <- HSD.test(ANOVA_RSR, trt = "Genotype", alpha = 0.05, console = TRUE)

\# Data framing to download

HSD_group_RSR <- HSD_result_RSR$groups

\# Save the data frame as CSV file

write.csv(HSD_group_RSR, file = "RSR_HSD_result.csv")

\## ANOVA: Specific root length (SRL)

ANOVA_SRL <- aov(SRL ~ Genotype, data = FinalDayData)

\## Tukey's HSD post-hoc test: specific root length (SRL)

HSD_result_SRL <- HSD.test(ANOVA_SRL, trt = "Genotype", alpha = 0.05, console = TRUE)

\# Data framing to download

HSD_group_SRL <- HSD_result_SRL$groups

\# Save the data frame as CSV file

write.csv(HSD_group_SRL, file = "SRL_HSD_result.csv")

\## ANOVA: primary root length (PRL)

ANOVA_PRL <- aov(PRL ~ Genotype, data = FinalDayData)

\## Tukey's HSD post-hoc test: primary root length (PRL)

HSD_result_PRL <- HSD.test(ANOVA_PRL, trt = "Genotype", alpha = 0.05, console = TRUE)

\# Data framing to download

HSD_group_PRL <- HSD_result_PRL$groups

\# Save the data frame as CSV file

write.csv(HSD_group_PRL, file = "PRL_HSD_result.csv")

\## ANOVA: first pair seminal root length (FPSRL)

ANOVA_FPSRL <- aov(FPSRL ~ Genotype, data = FinalDayData)

\## Tukey's HSD post-hoc test: first pair seminal root length (FPSRL)

HSD_result_FPSRL <- HSD.test(ANOVA_FPSRL, trt = "Genotype", alpha = 0.05, console = TRUE)

\# Data framing to download

HSD_group_FPSRL <- HSD_result_FPSRL$groups

\# Save the data frame as CSV file

write.csv(HSD_group_FPSRL, file = "FPSRL_HSD_result.csv")

\## ANOVA: second pair seminal root length (SPSRL)

ANOVA_SPSRL <- aov(SPSRL ~ Genotype, data = FinalDayData)

\## Tukey's HSD post-hoc test: second pair seminal root length (SPSRL)

HSD_result_SPSRL <- HSD.test(ANOVA_SPSRL, trt = "Genotype", alpha = 0.05, console = TRUE)

\# Data framing to download

HSD_group_SPSRL <- HSD_result_SPSRL$groups

\# Save the data frame as CSV file

write.csv(HSD_group_SPSRL, file = "SPSRL_HSD_result.csv")

\## ANOVA: root system width (RSW)

ANOVA_RW <- aov(RW ~ Genotype, data = FinalDayData)

\## Tukey's HSD post-hoc test: root system width (RSW)

HSD_result_RW <- HSD.test(ANOVA_RW, trt = "Genotype", alpha = 0.05, console = TRUE)

\# Data framing to download

HSD_group_RW <- HSD_result_RW$groups

\# Save the data frame as CSV file

write.csv(HSD_group_RW, file = "RW_HSD_result.csv")

\## ANOVA: convexhull area (CHA)

ANOVA_CHA <- aov(CHA ~ Genotype, data = FinalDayData)

\## Tukey's HSD post-hoc test: convexhull area (CHA)

HSD_result_CHA <- HSD.test(ANOVA_CHA, trt = "Genotype", alpha = 0.05, console = TRUE)

\# Data framing to download

HSD_group_CHA <- HSD_result_CHA$groups

\# Save the data frame as CSV file

write.csv(HSD_group_CHA, file = "CHA_HSD_result.csv")

Script number 9.

Tukeys HSD post hoc test for different root and shoot traits of different genotypes at day eight in high temperature stress condition.

\## Install and load the necessary packageslibrary(agricolae)

library(dplyr)

library(ggplot2)

library(broom)

library(tidyverse)

\# Memory cleaning and folder preparation

\# Clean up the system brain if any

rm(list=ls())

\# Select csv file by the following command

setwd("D:/251006_R_analysis")

FinalDayData_all <-read.csv("D:/251006_R_analysis/251230_FinalDayData_all_Genotype.csv")

\# Create a sub-set data for Control condition

FinalDayData <- subset(FinalDayData_all, Genotype %in% c("N61_H", "MSD034_H", "MSD054_H", "MSD296_H", "MSD392_H", "MSD417_H"))

Variables <- c("RD", "SH", "RDW", "SDW", "RSR","TRL", "SRL", "PRL", "FPSRL", "SPSRL",

"RW", "CHA", "FPSRA", "SPSRA")

\## ANOVA: rooting depth (RD)

ANOVA_RD <- aov(RD ~ Genotype, data = FinalDayData)

\## Tukey's HSD post-hoc test: rooting depth (RD)

HSD_result_RD <- HSD.test(ANOVA_RD, trt = "Genotype", alpha = 0.05, console = TRUE)

\# Data framing to download

HSD_group_RD <- HSD_result_RD$groups

\# Save the data frame as CSV file

write.csv(HSD_group_RD, file = "RD_HSD_result.csv")

\## ANOVA: shoot height (SH)

ANOVA_SH <- aov(SH ~ Genotype, data = FinalDayData)

\## Tukey's HSD post-hoc test: shoot height (SH)

HSD_result_SH <- HSD.test(ANOVA_SH, trt = "Genotype", alpha = 0.05, console = TRUE)

\# Data framing to download

HSD_group_SH <- HSD_result_SH$groups

\# Save the data frame as CSV file

write.csv(HSD_group_SH, file = "SH_HSD_result.csv")

\## ANOVA: root dry weight (RDW)

ANOVA_RDW <- aov(RDW ~ Genotype, data = FinalDayData)

\## Tukey's HSD post-hoc test: root dry weight (RDW)

HSD_result_RDW <- HSD.test(ANOVA_RDW, trt = "Genotype", alpha = 0.05, console = TRUE)

\# Data framing to download

HSD_group_RDW <- HSD_result_RDW$groups

\# Save the data frame as CSV file

write.csv(HSD_group_RDW, file = "RDW_HSD_result.csv")

\## ANOVA: shoot dry weight (SDW)

ANOVA_SDW <- aov(SDW ~ Genotype, data = FinalDayData)

\## Tukey's HSD post-hoc test: shoot dry weight (SDW)

HSD_result_SDW <- HSD.test(ANOVA_SDW, trt = "Genotype", alpha = 0.05, console = TRUE)

\# Data framing to download

HSD_group_SDW <- HSD_result_SDW$groups

\# Save the data frame as CSV file

write.csv(HSD_group_SDW, file = "SDW_HSD_result.csv")

\## ANOVA: root to shoot ratio (RSR)

ANOVA_RSR <- aov(RSR ~ Genotype, data = FinalDayData)

\## Tukey's HSD post-hoc test: root to shoot ratio (RSR)

HSD_result_RSR <- HSD.test(ANOVA_RSR, trt = "Genotype", alpha = 0.05, console = TRUE)

\# Data framing to download

HSD_group_RSR <- HSD_result_RSR$groups

\# Save the data frame as CSV file

write.csv(HSD_group_RSR, file = "RSR_HSD_result.csv")

\## ANOVA: Specific root length (SRL)

ANOVA_SRL <- aov(SRL ~ Genotype, data = FinalDayData)

\## Tukey's HSD post-hoc test: specific root length (SRL)

HSD_result_SRL <- HSD.test(ANOVA_SRL, trt = "Genotype", alpha = 0.05, console = TRUE)

\# Data framing to download

HSD_group_SRL <- HSD_result_SRL$groups

\# Save the data frame as CSV file

write.csv(HSD_group_SRL, file = "SRL_HSD_result.csv")

\## ANOVA: primary root length (PRL)

ANOVA_PRL <- aov(PRL ~ Genotype, data = FinalDayData)

\## Tukey's HSD post-hoc test: primary root length (PRL)

HSD_result_PRL <- HSD.test(ANOVA_PRL, trt = "Genotype", alpha = 0.05, console = TRUE)

\# Data framing to download

HSD_group_PRL <- HSD_result_PRL$groups

\# Save the data frame as CSV file

write.csv(HSD_group_PRL, file = "PRL_HSD_result.csv")

\## ANOVA: first pair seminal root length (FPSRL)

ANOVA_FPSRL <- aov(FPSRL ~ Genotype, data = FinalDayData)

\## Tukey's HSD post-hoc test: first pair seminal root length (FPSRL)

HSD_result_FPSRL <- HSD.test(ANOVA_FPSRL, trt = "Genotype", alpha = 0.05, console = TRUE)

\# Data framing to download

HSD_group_FPSRL <- HSD_result_FPSRL$groups

\# Save the data frame as CSV file

write.csv(HSD_group_FPSRL, file = "FPSRL_HSD_result.csv")

\## ANOVA: second pair seminal root length (SPSRL)

ANOVA_SPSRL <- aov(SPSRL ~ Genotype, data = FinalDayData)

\## Tukey's HSD post-hoc test: second pair seminal root length (SPSRL)

HSD_result_SPSRL <- HSD.test(ANOVA_SPSRL, trt = "Genotype", alpha = 0.05, console = TRUE)

\# Data framing to download

HSD_group_SPSRL <- HSD_result_SPSRL$groups

\# Save the data frame as CSV file

write.csv(HSD_group_SPSRL, file = "SPSRL_HSD_result.csv")

\## ANOVA: root system width (RSW)

ANOVA_RW <- aov(RW ~ Genotype, data = FinalDayData)

\## Tukey's HSD post-hoc test: root system width (RSW)

HSD_result_RW <- HSD.test(ANOVA_RW, trt = "Genotype", alpha = 0.05, console = TRUE)

\# Data framing to download

HSD_group_RW <- HSD_result_RW$groups

\# Save the data frame as CSV file

write.csv(HSD_group_RW, file = "RW_HSD_result.csv")

\## ANOVA: convexhull area (CHA)

ANOVA_CHA <- aov(CHA ~ Genotype, data = FinalDayData)

\## Tukey's HSD post-hoc test: convexhull area (CHA)

HSD_result_CHA <- HSD.test(ANOVA_CHA, trt = "Genotype", alpha = 0.05, console = TRUE)

\# Data framing to download

HSD_group_CHA <- HSD_result_CHA$groups

\# Save the data frame as CSV file

write.csv(HSD_group_CHA, file = "CHA_HSD_result.csv")
